# The K18-hACE2 Transgenic Mouse Model Recapitulates Non-Severe and Severe COVID-19 in Response to Infectious Dose of SARS-CoV-2 Virus

**DOI:** 10.1101/2021.05.08.443244

**Authors:** Wenjuan Dong, Heather Mead, Sierra Jaramillo, Tasha Barr, Daniel S. Kollath, Vanessa K. Coyne, Nathan E. Stone, Ashley Jones, Jianying Zhang, Aimin Li, Li-Shu Wang, Martha Milanes-Yearsley, Paul S Keim, Bridget Marie Barker, Michael Caligiuri, Jianhua Yu

## Abstract

A comprehensive analysis and characterization of a SARS-CoV-2 infection model that mimics non-severe and severe COVID-19 in humans is warranted for understating the virus and developing preventive and therapeutic agents. Here, we characterized the K18-hACE2 mouse model expressing human (h)ACE2 in mice, controlled by the human keratin 18 (K18) promoter, in epithelia, including airway epithelial cells where SARS-CoV-2 infections typically start. We found that intranasal inoculation with higher viral doses (2×10^3^ and 2×10^4^ PFU) of SARS-CoV-2 caused lethality of all mice and severe damage of various organs, including lungs, liver, and kidney, while lower doses (2×10^1^ and 2×10^2^ PFU) led to less severe tissue damage and some mice recovered from the infection. In this humanized hACE2 mouse model, SARS-CoV-2 infection damaged multiple tissues, with a dose-dependent effect in most tissues. Similar damage was observed in biopsy samples from COVID-19 patients. Finally, the mice that recovered after infection with a low dose of virus also survived rechallenge with a high dose of virus. Compared to other existing models, the K18-hACE2 model seems to be the most sensitive COVID-19 model reported to date. Our work expands the information available about this model to include analysis of multiple infectious doses and various tissues with comparison to human biopsy samples from COVID-19 patients. In conclusion, the K18-hACE2 mouse model recapitulates both severe and non-severe COVID-19 in humans and can provide insight into disease progression and the efficacy of therapeutics for preventing or treating COVID-19.

**Importance:** The pandemic of COVID-19 has reached 112,589,814 cases and caused 2,493,795 deaths worldwide as of February 23, 2021, has raised an urgent need for development of novel drugs and therapeutics to prevent the spread and pathogenesis of SARS-CoV-2. To achieve this goal, an animal model that recapitulates the features of human COVID-19 disease progress and pathogenesis is greatly needed. In this study, we have comprehensively characterized a mouse model of SARS-CoV-2 infection using K18-hACE2 transgenic mice. We infected the mice with low and high doses of SARS-CoV-2 virus to study the pathogenesis and survival in response to different infection patterns. Moreover, we compared the pathogenesis of the K18-hACE2 transgenic mice with that of the COVID-19 patients to show that this model could be a useful tool for the development of anti-viral drugs and therapeutics.

## Introduction

The global pandemic of coronavirus disease 2019 (COVID-19) caused by the extremely contagious RNA coronavirus SARS-CoV-2 has led to 112,589,814 cases and 2,493,795 deaths worldwide as of February 23, 2021 (1). The median time to development of symptoms is 5.1 days after exposure to SARS-CoV-2 (2). The median time from onset to clinical recovery for mild cases is approximately 2 weeks, with 3-6 weeks for patients with severe or advanced disease (3). The main symptoms, such as fever or chills, cough, shortness of breath, difficulty breathing, and sore throat normally ease after recovery; however, 3-17% patients, especially the elderly and individuals with cancer and diabetes, develop rapid viral replication and severe lung damage, resulting in severe disease with greatly increased risk of death (4). Many groups are trying to understand this rapid disease progression and studies suggest that it may involve an impaired immune response (5-7). A mouse model that appropriately mirrors progression of the human disease should help the development of vaccines or therapeutics for COVID-19 and could be used as a conceptual basis for rapid response to future viral pandemics.

During infection with SARS-CoV-2, the coronavirus spike (S) glycoprotein promotes SARS-CoV-2 entry into host cells via the host receptor angiotensin converting enzyme 2 (ACE2)(8). K18-hACE2 transgenic mice, which were originally developed to study the infection of SARS-CoV, express human ACE2, the receptor used by SARS-CoV-2 to gain entry into cells (9). The human keratin 18 (K18) promoter is responsible for directing expression to airway epithelial cells, where respiratory infections usually begin. Recent research reported this SARS-CoV-2 hACE2 mouse infection model develops severe interstitial pneumonia with high viral loads into the lungs and immune cell infiltration into the alveoli (10-13). A comprehensive analysis and characterization of this model will be useful for studying the mechanisms of pathogenesis by SARS-CoV-2 and developing preventive, therapeutic, and vaccinal agents.

In the current study, we evaluate K18-hACE2 transgenic mice in response to multiple infectious doses, using multiple tissues, to gain a complete understanding of the response to the infectious dose. We also evaluated the response of K18-hACE2 mice to high-dose re-infection after recovery from low-dose infection.

## Results

### The K18-hACE2 model is a lethal infection model for SARS-CoV-2 infection

To understand the effects of SARS-CoV-2 viral dosage on pathogenesis and survival, we established an infection model using the K18-hACE2 transgenic mice, in which human angiotensin-converting enzyme 2 (hACE2) expression is driven by the epithelial cell-specific promoter K18 in C57BL/6 mice(9). The mice were infected with SARS-CoV-2 at 2 ×10^1^ PFU (low dose), 2 ×10^2^ PFU (low dose), 2 ×10^3^ PFU (high dose), 2 ×10^4^ PFU (high dose) or PBS vehicle control via intranasal inoculation. Blood was collected at day 3 post infection (p.i.) then the mice were euthanized, and viral loads and pathology were examined at day 6 p.i. (Fig. 1A). We found that mouse body weight decreased in mice infected with 2 ×10^3^ PFU and 2 ×10^4^ PFU by day 4 p.i, whereas mice infected with 2 ×10^1^ PFU and 2 ×10^2^ PFU showed a less dramatic body weight decrease; a body weight decrease of 20% occurred at days 5–6 in the higher-dose groups, but took until day 10–11 for 80% of the mice in the low-dose groups (**Fig. 1B**). 30% of the mice from the lower-dose groups even started to gain weight again at day 4 (for 2×10^1^) and day 8 (for 2×10^2^) (**Fig. 1B**), indicating a recovery from the viral infection in low-dose groups. Consistent with body weight data, 90% of the mice infected with 2 ×10^3^ PFU or 2 ×10^4^ PFU died at approximately at day 7, whereas 50% of the mice infected with 2 ×10^1^ PFU or 2 ×10^2^ PFU died approximately at day 10 (**Fig. 1C**). Three mice infected with 2 ×10^1^ PFU or 2 ×10^2^ PFU, and none infected with 2 ×10^3^ PFU or 2 ×10^4^ PFU, recovered and survived until the end of follow-up on day 20 p.i. (**Fig.1C**).

**Figure 1.**
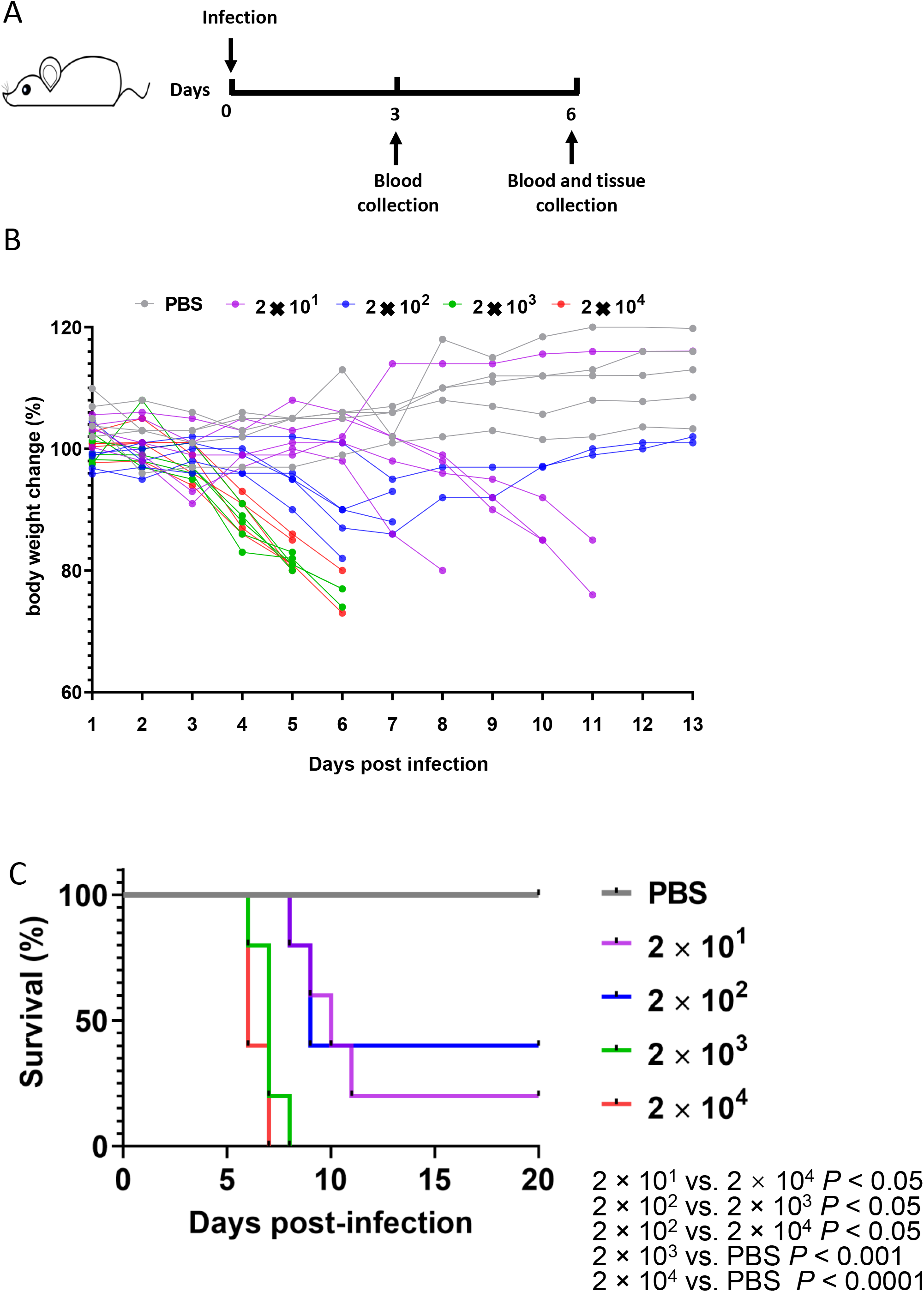
K18-hACE2 mouse infection model with high and low dose of SARS-CoV-2. A. Experimental scheme of the K18-hACE2 mouse infection model. The mice were intranasally infected with 2 ×10^1^ PFU, 2 ×10^2^ PFU, 2 ×10^3^ PFU or 2 ×10^4^ PFU virus. Blood samples were collected at 3- and 6-days post infection. Tissue samples were collected at 6 days post infection. Mouse body weights (B) and survival (C) were monitored daily for 13 days. Each dot represents one mouse at the indicated time point.

### Expression distribution of viral genes in the tissues of K18-hACE2 mice infected with high or low doses of SARS-CoV-2

To further investigate the viral infection pattern, we used reverse transcriptase (RT)-PCR to measure viral spike protein RNA levels in the brain, trachea, lung, liver, spleen, small intestine, stomach, large intestine, kidney, and testis. The mean viral loads in all three higher dose groups were over PBS and 1 × 10^2^, except for 2 ×10^1^ PFU in the spleen, and no virus was detected in uninfected controls (**Fig. 2A**). We observed three types of dose-response relationships. In trachea, lung, stomach, and kidney, there was a stepwise dose-response relationship. In heart, liver, spleen, and small intestine, the observed a plateau, with the 3 higher dose groups having similar viral loads. In brain, large intestine, and testis, mice infected with 2 ×10^1^ PFU or 2 ×10^2^ PFU had similar viral loads and mice infected with 2 ×10^3^ PFU or 2 ×10^4^ PFU had much higher viral loads. This indicates that there are tissue-specific factors modulating the effect of dose on viral copy number. The viral RNA levels were high in the lungs of animals in all dose groups (**Fig. 2A**), consistent with the notion the lung is an important site of infection, especially at lower viral doses. The viral RNA levels were also very high in brain for mice infected with 2 ×10^3^ PFU or 2 ×10^4^ PFU. We used immunohistochemistry (IHC) to measure expression of SARS-CoV-2 nucleocapsid protein (NP) in formalin-fixed, paraffin-embedded (FFPE) tissues, which confirmed the infection pattern in the lungs and brains of the K18-hACE2 mice infected with a series of increasing viral doses, demonstrating that viral spread in tissues was also dose-dependent (**Fig. 2B**).

**Figure 2.**
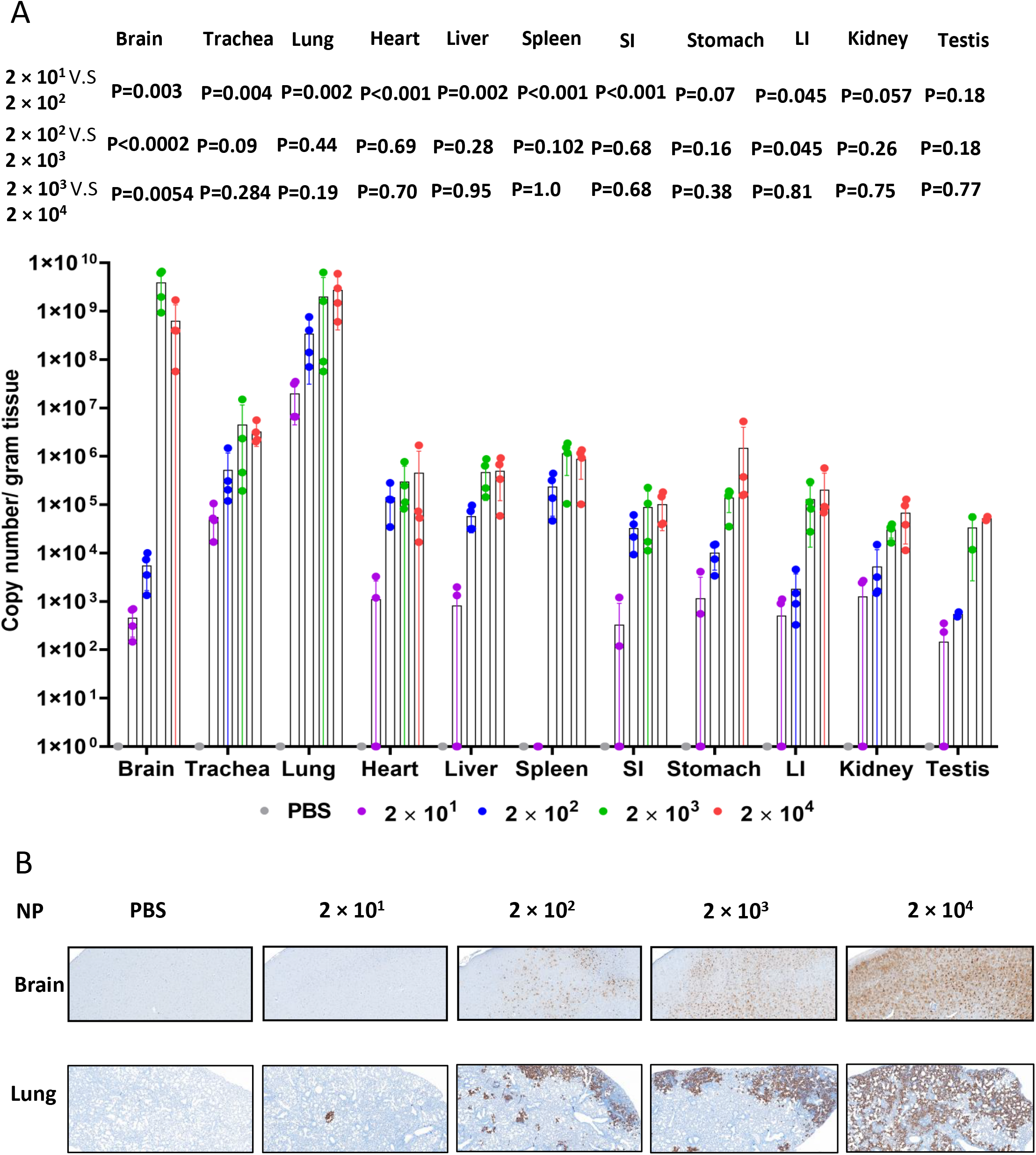
Viral quantification in mice after SARS-CoV-2 infection. A. Viral RNA levels are shown for brain, trachea, lung, heart, liver, spleen, small intestine (SI), stomach, large intestine (LI), kidney and testis. B. Viral nucleocapsid protein (NP) was detected in brain and lung of the mice infected with high and low dose of SARS-CoV-2 (Scale bar, 40µm).

To evaluate the lung cell types that are susceptible to SARS-CoV-2 infection, we performed IHC on samples from mice infected with 2 × 10^4^ PFU. We double stained lung sections for viral NP and CCL10, which is the marker of club (clara) cells, or SPC, the marker of alveolar type 2 cells, respectively. We found that only few club cells or alveolar type 2 cells co-localized with NP, suggesting they are susceptible to SARS-CoV-2 infection (**Fig. 3A**). Macrophages are a major immune cell population in lung, where they act as a first-line defense against invading pathogens. We therefore used the CD68 as a macrophage marker to examine if SARS-CoV-2 could infect tissue resident macrophages in lung. We noticed that that CD68 positive cells increased dramatically in response to SARS-CoV-2 infection, although very few macrophages stained positive for NP, suggesting that it could be possible for SARS-CoV-2 to suppress immune response via infection of lung macrophages (**Fig. 3A**). Consistent with the finding in the K18-hACE2 mouse model, in biopsy lung samples from COVID-19 patients, few viral spike protein is colocalized with alveolar type 2 cells and CD68-positive cells (**Fig. 3B**). These results are in line with another SARS-CoV-2 infection mouse model that uses CRISPR/Cas9 knockin of *hACE2* into Exon 2 of the *mAce2* gene, as well as COVID-19 patient samples, which show SARS-CoV-2 in macrophages of pulmonary alveolus (12, 14). Given the high levels of SARS-CoV-2 in brain, we examined whether neurons are infected with SARS-CoV-2. We found strong overlap between staining for the neuronal marker Neu and staining for NP (**Fig. 3A**). Human lung alveolar type II cells were injured significantly by COV19 virus infection. This was identified by the reduction of the number and stain intensity of SPC positive cells (yellow) in infected lung compared to no-infected lung. In COVID-19 lung sample #1, which has highest virus amount (teal), the SPC positive cells are almost completely lost, whereas in the other lungs, SPC positive cells are low and fallen off from epithelium into alveolar lumen; meanwhile, Consistent with findings in mouse COV19 model, in human lung, very few CD68 positive macrophages (yellow) are also positive for Spike (teal). Our results indicated that the K18-hACE2 mice could be a suitable model for mimicking the infections of the COVID-19 patients.

**Figure 3.**
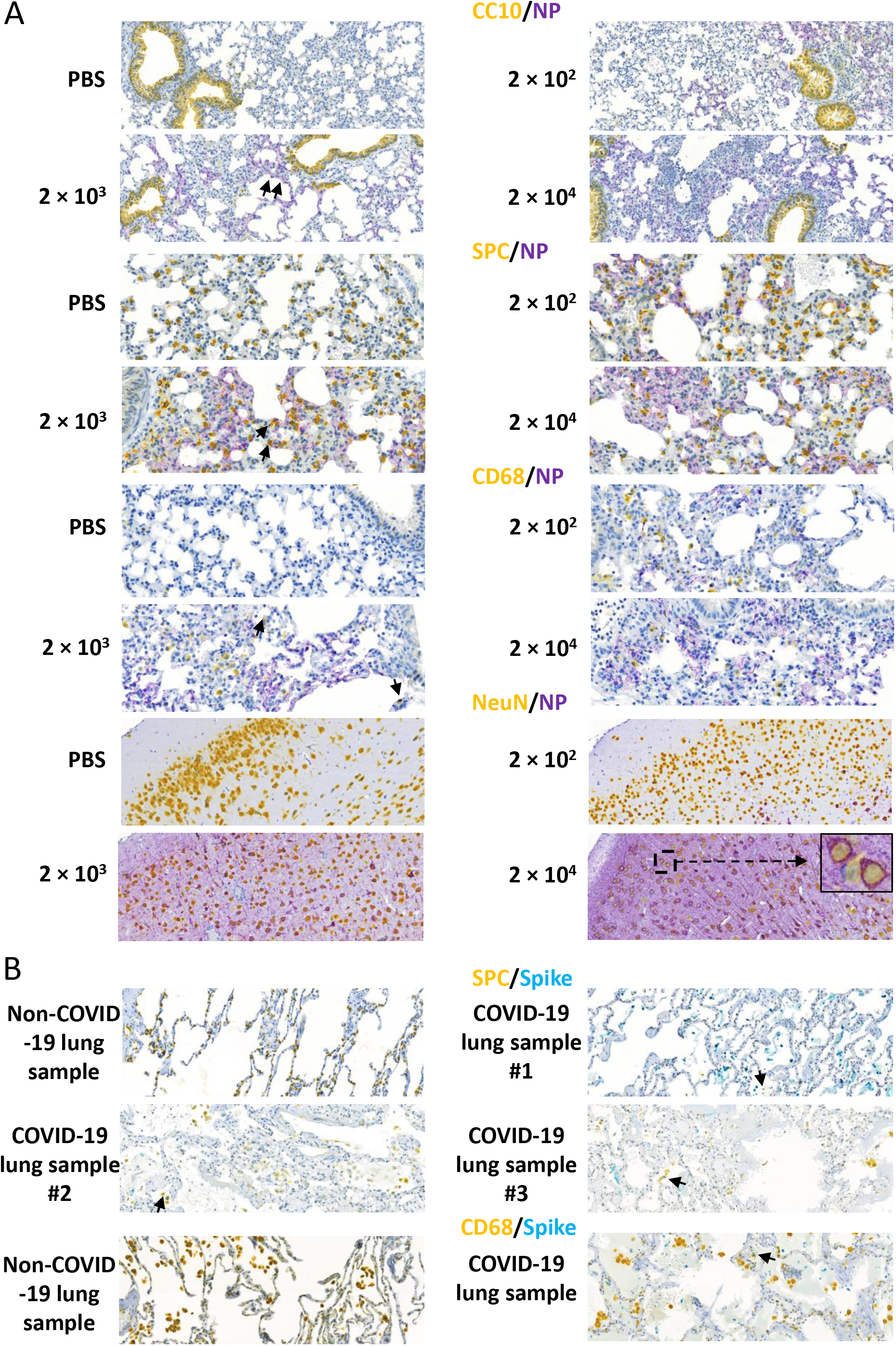
Viral distribution in mice after SARS-CoV-2 infection. A. Representative images show double staining of NP (purple) with lung club (Clara) cells and alveolar type 2 cells using the markers CCL10 (yellow) and SPC (yellow), macrophages using the CD68 marker (yellow) (20 × magnification), and neurons cells using the NeuN marker (yellow), in mice infected with high-dose SARS-CoV-2 (10 × magnification, with 40 × magnification in the 2 ×10^4^ PFU group). Black arrows indicate double staining cells. B. Representative images show double staining of spike protein (teal) with alveolar type 2 cells using SPC (yellow) as a marker and macrophages using CD68 (yellow) as a marker in COVID-19 patient samples (20 × magnification). Black arrows indicate double staining cells.

### Pathology of tissues of K18-hACE2 mice infected with SARS-CoV-2

To assess disease severity, we used H&E staining to assess pathology in multiple FFPE tissues from K18-hACE2 mice infected with a series of increasing doses of SARS-CoV-2. As expected, lung tissues showed the most severe damage. Mice in the low-dose groups showed an average of 30–60% more alveolar congestion and consolidation compared to uninfected mice (**Fig. 4A**). Some lung tissue showed alveolar hemorrhage as well as lymphocytic pneumonitis with alveolar thickening and peripheral parenchymal collapse. Lung damage in the high-dose groups were even more extensive, consisting of ∼20% alveolar collapse with ruptured septa, as well as thickened alveolar septa and intra-alveolar. Parenchymal consolidation was evident in 50% of the tissue, along with interstitial inflammation and pneumonitis (**Fig. 4A**). Similarly, trachea tissues showed mild epithelial damage in the lower-dose groups and more severe epithelial damage in the higher-dose groups. Consistent with the brain showing high viral gene expression for the higher-doses groups (**Fig. 2A**), brain tissues showed mild congestion in some mice from the low-dose groups, with more extensive congestion in the high-dose groups. In liver tissue, the low-dose groups showed several foci of spotty and patchy necrosis (focal perivenular), Kupffer cell hyperplasia, and focal portal inflammation; the high-dose groups also showed Kupffer cell hyperplasia along with congestion, reactive change, apoptotic hepatocytes, and focal lobular inflammation (**Fig. 4A**). In spleen, the low dose group showed red pulp congestion while the high dose group also showed lymphoid hyperplasia and mild extramedullary hematopoiesis. The stomach and small and large intestines in the low-dose groups appeared normal but the stomach showed reactive changes, diffuse epithelial sloughing and chronic inflammation along the myenteric plexus, while the small and large intestine showed myenteric plexus inflammation. The testis in the low dose group also appeared normal but showed focal tubular damage and congestion in the high dose group. In kidney, both the low dose and high dose group showed cortical congestion, tubular damage and focal tubular collapse. Meanwhile, both low dose and high dose group showed ischemic change and disarray in heart (**Fig. 4A**). These findings indicate that SARS-CoV-2 infection damages multiple tissues, with a dose-dependent effect in most tissues.

**Figure 4.**
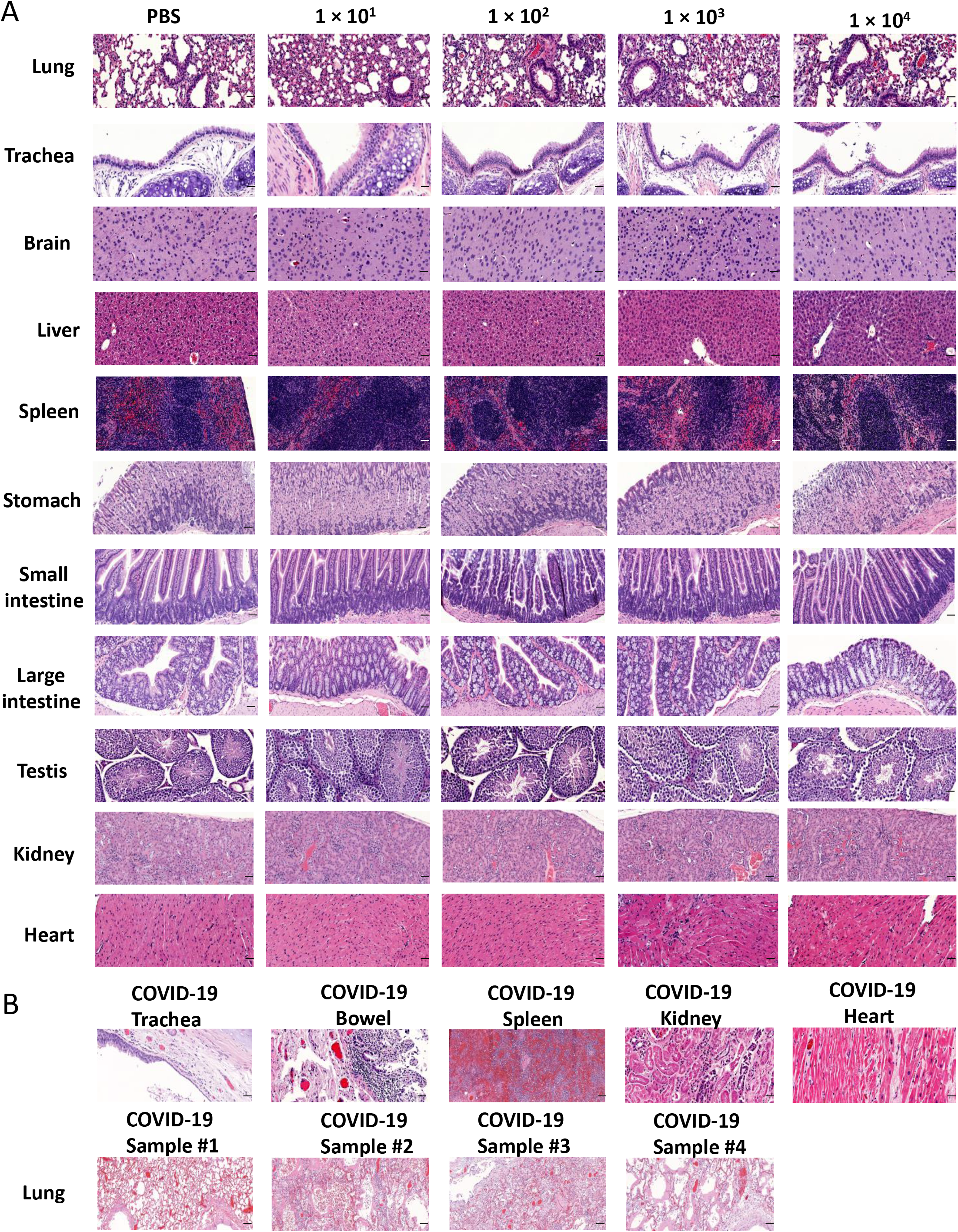
Pathological changes in multiple tissues of K18-hACE2 mice infected with the indicated dose of SARS-CoV-2 or autopsy tissue from COVID-19 patients. A. Tissue damage in brain, trachea, lung, heart, liver, spleen, small intestine, stomach, large intestine, kidney and testis of K18-ACE2 mice after SARS-CoV-2 infection (Scale bar, 40µm). B. Tissue damage in trachea, bowel, spleen, kidney, heart, and lung in COVID-19 patient samples (Scale bar, 40µm).

To compare the pathology in the mouse model to COVID-19 patients, we also examined the pathology in autopsy samples of COVID-19 patients. We identified congestion and inflammation in multiple organs, as well as the epithelial damage and necrosis, which was consistent with our findings in the SARS-CoV-2-infected K18-hACE2 mouse model (**Fig. 4B**).

### Expression of hACE2 in K18-hACE2 mice and human tissues

To better understand the infection pattern and virus distribution in this mouse model, we measured hACE2 RNA expression levels in various tissues of hACE2 mice by RT-PCR. We found that, compared to the expression of hACE2 in lung tissue, there was also high expression in brain, trachea, heart, stomach, small intestine, large intestine, kidney, and testis; whereas there was lower expression in the liver and spleen (**Fig. 5A**). We also confirmed the hACE2 expression in brain, trachea, lung and kidney of the k18-hACE2 mice at the protein level by IHC (**Fig. 5B**). The expression of hACE2 in these three tissues in humans is consistent with the finding that they had the highest viral load, further confirming the K18-hACE2 mouse is a good model for studying in vivo SARS-CoV-2 infection. We also examined hACE2 expression in a human tissue array. We found that hACE2 is not only highly expressed in lung but also in stomach, small and large intestine, kidney and testis. However, the expression of ACE2 in human brain was not as high as in the brain of K18-hACE2 mice; this could be due to the insertion of the K18 promotor, which confers higher expression of hACE2 in the mouse model (**Fig. 5C**). This finding is consistent with another transgene driven by a human K18 regulatory element in mice(15).

**Figure 5.**
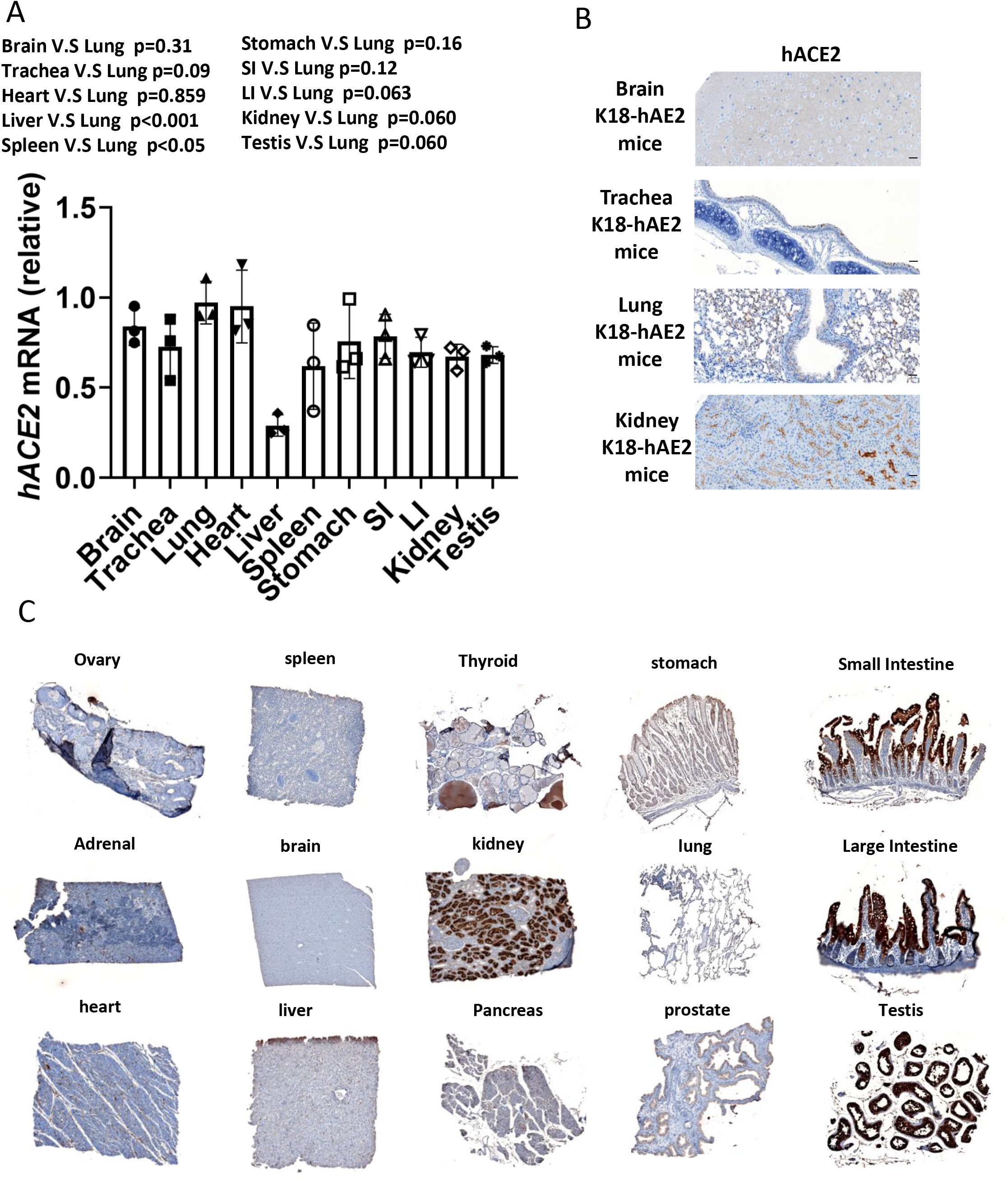
Tissue distribution of hACE2 in K18-hACE2 mice and human samples. A. Detection of NP RNA in multiple tissues of K18-hACE2 mice by RT-qPCR. B. hACE2 expression in brain, trachea, lung and kidney of K18-hACE2 mice by IHC. C. Human tissue array IHC staining for hACE2 protein.

### Protective role of serum from previously infected mice and the mice with previous low-dose of virus infection against rechallenge with high-dose of virus infection in K18-hACE2 mice

To evaluate if the serum from infected mice could protect the mice from viral infection, we injected virus-naïve mice with serum from mice infected with 2 × 10^4^ PFU group of mice and then infected the virus-naïve mice with SARS-COV2 at 2 × 10^4^ PFU/mouse. The body weights of treated vs. control mice were continuously monitored. We found that the mice treated with serum from infected mice showed a delayed and slower body weight drop compared to the untreated group (**Fig. 6A**), suggesting the serum from infected mice could have a protective effect on newly infected mice. Serum from infected mice did not significantly protect but show but show a trend against death after virus rechallenge, which might be due to a little amount of serum that we can collected to have enough sample size (**Fig. 6B**).

**Figure 6.**
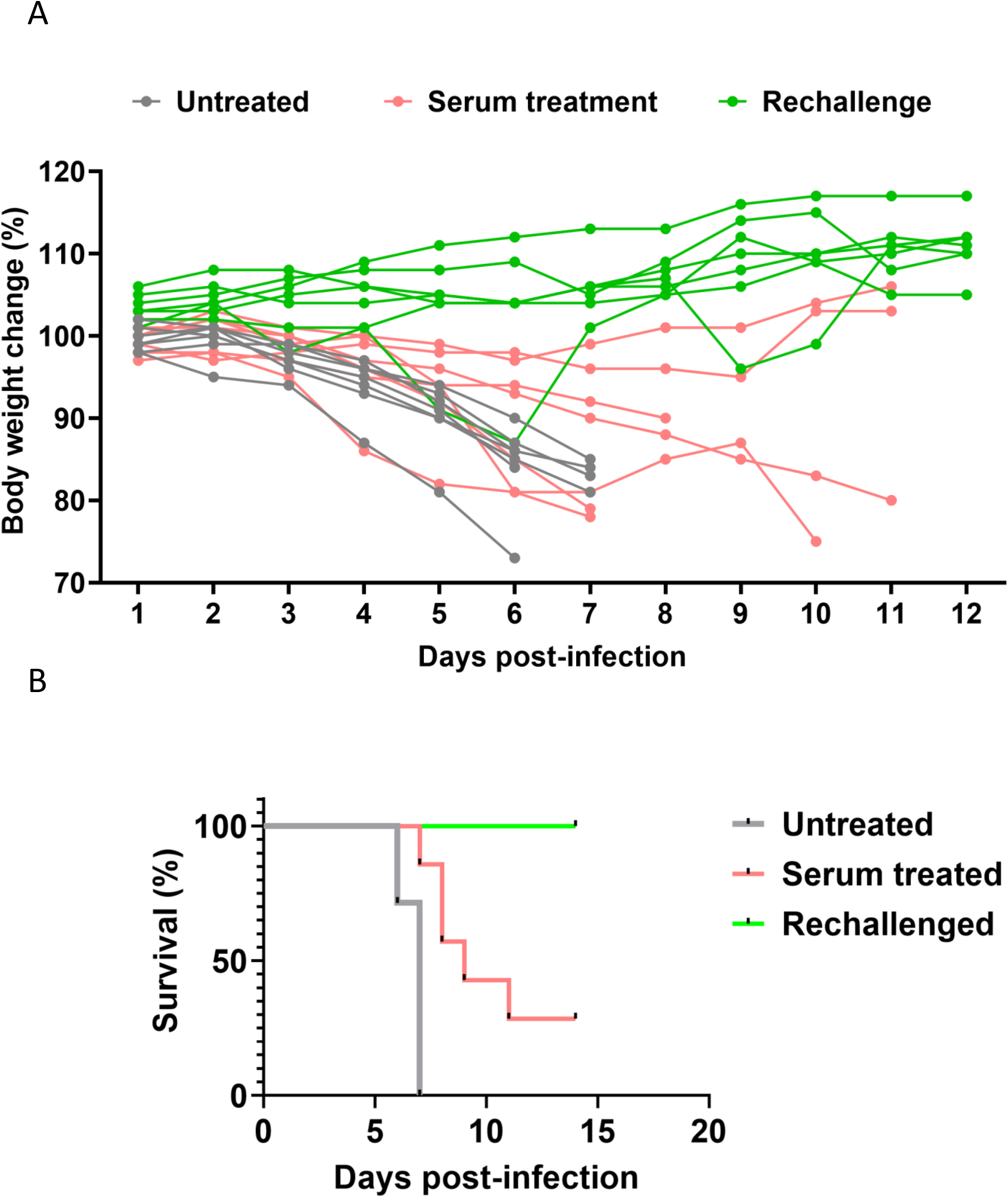
Protective role of serum from previously infected mice and same mice with previous infection against rechallenge in K18-hACE2 mice. A. Body weight of mice treated with serum from previously infected mice and same mice with previously low dose-infected mice after challenge with a high dose of SARS-CoV-2. B. Survival of mice treated with serum from previously low dose-infected mice and same mice with previous infection after challenge with a high dose of SARS-CoV-2. Mice with serum protection were infected with 2 ×10^4^ PFU SASR-CoV-2 24 h after being infused with serum from mice infected with 2 ×10^4^ PFU virus for 6 days. For the rechallenge, mice were infected with 2 ×10^1^ PFU or 2 ×10^2^ PFU SASR-CoV-2 for two weeks and rechallenge with 2 ×10^4^ PFU SASR-CoV-2.

In addition, we rechallenged the mice that survived 2 × 10^1^ or 2 × 10^2^ PFU infection with 2 × 10^4^ PFU virus when they recovered to the initial body weight. We found that all mice survived the high-dose virus re-challenge and did not show substantial body weight drop, suggesting that recovery from low-dose infection conferred anti-viral activity that protected against rechallenge with high dose virus (**Fig. 6B**).

## Discussion

The recent outbreak of SARS-CoV-2 that has claimed the lives of nearly 2,500,000 people has led to an urgent need for a mouse infection model that accurately mirrors inoculation, development and progression of the human disease termed COVID-19. The K18-hACE2 model was developed to study coronavirus infection (9).In the current study, we performed a comprehensive analysis of viral load and tissue pathology based on a range of inoculation doses and compared the results to observations in biopsy samples from COVID-19 patients. We found a dose-response relationship between infectious dose and loss of body weight, viral titer, tissue pathology and mortality. Notably, there was not a uniform relationship between infectious dose and viral titer, with some tissues accumulating virus at lower doses and others accumulating virus at higher doses. We evaluated hACE2 levels in the K18-ACE2 mice and found similarity to ACE2 levels in humans. We also showed that challenge with a low dose of SARS-CoV-2 protected against the lethality of rechallenge with a higher dose. Overall, our results indicate that low dose infection in the K18-hACE2 model mimics non-severe COVID-19, while infection with higher doses mimics severe COVID-19.

Due to the unprecedented impact of the COVID-19 pandemic, many models of human COVID-19 are under investigation. In addition to the K18-hACE2 model, other mouse models have been described. Hassan *et al*. used replication-defective adenoviruses encoding human ACE2-tranduced mice to infect BALB/c mice, followed by SARS-CoV-2 infection, to model COVID-19 (13). Following infection by 10^5^ focus-forming units (FFU) of SARS-CoV-2, the body weight of mice was maintained but did not drop, therefore it seems that the model is not a lethal model. Using CRISPR/Cas9 technology to knock-in hACE2, Sun *et al*. established a model (12). In response to 4×10^5^ PFU of SARS-CoV-2, the animals experienced robust viral replication in multiple tissues and interstitial pneumonia, but no obvious clinical symptoms or mortality; only 10% of aged mice lost their weight at day 3 p.i. and recovered. With the low incidence rate and very mild symptoms, this model is not ideal to recapitulate human COVID-19. This might be due to the nature of the knock-in with only one copy of hACE2 systemically existing in mice. Qin and colleagues developed a hACE2 transgenic mouse model with a mouse ACE2 promoter for SARS and used the model to study SARS-CoV-2 infection (11, 16). Infected by 10^5^ TCID50 of SARS-CoV-2, the mouse body weights dropped at day 1 and continued until day 5, and then almost completely recovered on day 14 without lethality, suggesting that this model at least does not mimic severe COVID-19 patients. This might be due to lower promoter activity or gene expression of murine (m)ACE2 compared to the K18 promoter (16). Ostrowski *et al*. developed transgenic mice using the promoter of the human FOXJ1, a transcription factor required for differentiation of ciliated epithelial cells in the airway, to drive hACE2 expression (17). Jiang and colleagues used this model to study SARS-CoV-2 (10). Infected with 3×10^4^ PFU virus, among 10 mice, 2 had no infection, 4 had no drop in body weight, and 4 had body weight drops with 3 deaths and 1 recovery (10). In contrast, for the K18-hACE2 model that we characterized here, with the dose of 2×10^3^ PFU, which is lower than the doses used in all above mouse models, all mice succumbed to infection with body weight drops and mortality of 100% by day 8. With lower dose infection such as 2×10^1^ and 2×10^2^ PFU, 30% of the mice recovered. Collectively, these suggest that the K18-hACE2 model is the most sensitive model for COVID-19, with ability to recapitulate aspects of human disease from both non-severe and severe COVID-19 patients. Our results are supported by those from other groups (18-22).This might be due to unique aspects of the K18-hACE2 model: 1) hACE2 is used, where in contrast other models including golden hamster, ferret, cat, Chinese tree shrew and even mouse models with an endogenous ACE2 promoter recapitulate human expression levels (23). 2) Multiple copies of hACE2 are placed in the murine genome, where as one copy of hACE2 may not be ideal, evidenced by the knock-in study by Sun et al (12); and 3) human K18 is likely stronger than the promoters in other models, such as the human FOXJ1 in the study by Jiang et al. (10)

The pathological damage in lung and brain in the K18-hACE2 mouse infection model are similar to the clinical symptoms of COVID-19 patients, suggesting this mouse model can recapitulate SARS-CoV-2 infection in humans. However, the K18-hACE2 mouse model was originally generated by inserting hACE2 in the mouse genome under the human K18 promoter, which may not result in the exact distribution of ACE2 expression as in humans. In fact, we observed high expression levels of hACE2 in mouse brains, correlating with a higher virus titer in the brain compared to most other organs or tissues; however, in biopsy samples of humans, the hACE2 expression is very low. Nonetheless, brain infections with COVID-19 and deaths of patients from brain-specific COVID infection have been frequently reported (24-26). Interestingly, our study showed that SARS-CoV-2 can infect neurons in an animal model, which is consistent with previous studies using human neuron organoids (27-29). It will be interesting to know whether this neuronal infection causes taste and smell loss, commonly found in patients with COVID-19, as neuronal circuits respond to gustatory and olfactory cues (30).

In summary, we characterized the K18-hACE2 model for COVID-19 in comparison to human biopsy samples. The model shows dose-dependent sensitivity to SARS-CoV-2 infection. Infection with low doses recapitulates the disease observed in non-severe COVID-19 patients, while infection with higher doses recapitulates the disease observed in patients with severe COVID-19. The K18-hACE2 humanized COVID-19 mouse model is excellent to study COVID-19 and develop preventive and therapeutic drugs, as well as vaccines, for coronavirus diseases.

## Materials and Methods

### Ethics statement

Mouse-model studies were performed in an animal biosafety level 3 (ABSL3) facility. The animal protocol of these studies was reviewed and approved by the institutional animal care and use committee of Northern Arizona University (protocol #20-005).

### Viruses, cells and mice

The SARS-COV-2 strain used in the mouse model was SARS-CoV-2/human/USA/WA-CDC-WA1/2020, which was purchased from EBI. The viruses were amplified using Vero-E6 cells (ATCC). Vero-E6 cells were cultured in Dulbecco’s modified Eagle’s medium (DMEM) plus 10% fetal bovine serum (FBS) at 37□ and 5% CO_2_. The cells were inoculated with virus at a multiplicity of infection (MOI) of 0.001 and cultured for 96 h. Then the supernatant was collected and titrated using a plaque assay.

The K18-hACE2 transgenic mice, which use the human keratin 18 (KRT18) promoter to direct human ACE2 expression, were purchased from the Jackson Laboratory.

### Mouse infection

Female and Male of six-to-eight-week-old K18-hACE2 transgenic mice under the C57BL/6J background were anesthetized and intranasally (i.n.) infected with SARS-COV-2 virus at a dosage of 2 × 10^1^ PFU/mouse, 2 × 10^2^ PFU/mouse, 2 × 10^3^ PFU/mouse or 2 × 10^4^ PFU/mouse. The uninfected control mice were inoculated with phosphate-buffered saline (PBS). All the mice were observed and weighed daily. Blood was collected on day 6 from high dose groups and used for serum protection experiments. The mice were euthanized at day 6 post infection and the tissues were collected for further analysis.

### Viral rechallenge model

The mice that survived i.n. infection with 2 × 10^1^ PFU and 2 × 10^2^ PFU of virus were rechallenged two weeks after the initial infection with 2 × 10^4^ PFU virus i.n. when they recovered to the initial body weight after the first infection. The body weight was continuously monitored.

### Serum protection model

The serum of the mice infected by 2 × 10^4^ PFU virus was collected at day 6 post infection. Half of the serum was intraperitoneally injected into the non-infected mice 2 days before the mice were i.n. infected with 2 × 10^4^ PFU virus. The other half of the serum was i.n. administered right before the infection.

### Viral titer detection

The tissues were collected, weighed and immediately homogenized using an electric homogenizer (Thomas Scientific). After centrifugation at 1200g for 10 min, the supernatant was isolated and used for viral titer detection.

### Viral RNA copy number detection

The viral RNA was isolated from the homogenized tissues using the PureLink RNA Mini kit (Invitrogen). A one-step RT-PCR kit (BioRad) was used to detect the viral RNA using Applied Biosystems QuantStudio 12K Flex Real-Time PCR System with the following cycling protocol: reverse transcription at 50°C for 10 minutes; hot start at 95°C for 10 minutes; and 40 cycles of denaturation at 95°C for 10 seconds and annealing at 60°C for 30 seconds. The primer sequences were CoV2-S_19F (5’ -GCTGAACATGTCAACAACTC-3’) and CoV2-S_143R (5’ -GCAATGATGGATTGACTAGC-3’), which were designed to target a 125 bp region of the SARS-CoV-2 spike protein (31). The standard samples were serial 10-fold dilutions of a known copy number of the HKU1 virus. The results were normalized and expressed as genome equivalent copies per gram of tissue.

### Human ACE2 RNA quantification

Total RNA from the indicated tissues of the K18-hACE2 mice was isolated using the PureLink RNA Mini Kit (Invitrogen) and cDNA was synthesized using SuperScript Reverse Transcriptase (Thermo Fisher). Human ACE2 expression was examined using the following primers: F: CGAAGCCGAAGACCTGTTCTA; R: GGGCAAGTGTGGACTGTTCC, under the following PCR conditions: 98 °C for 30 s, followed by 40 cycles of 98 °C for 15 s, 62 °C for 30 s and 72 °C for 60 s.

### Immunohistochemistry

Tissues were harvested from the infected K18-hACE2 mice and immediately fixed in 10% neutral buffered formalin. Dehydration, clear and paraffinionization was performed on a Tissue - Tek VIP Vacuum Infiltration Processor (SAKURA). The samples were embedded in paraffin using a Tissue-Tek TEC Tissue Embedding Station (SAKURA). Samples were then sectioned at 5 μm and put on positively charged glass slides. The slides were deparaffinized, rehydrated and stained with Modified Mayer’s Hematoxylin and Eosin Y Stain (America MasterTech Scientific) on a H&E Auto Stainer (Prisma Plus Auto Stainer, SAKURA) according to standard laboratory procedures.

Single or double IHC stains were performed on Ventana Discovery Ultra (Ventana Medical Systems, Roche Diagnostics, Indianapolis, USA) IHC Auto Stainer. Briefly, the slides were loaded on the machine, deparaffinization, rehydration, endogenous peroxidase activity inhibition and antigen retrieval were first performed. For single IHC stain, the primary antibodies were incubated with DISCOVERY anti-Rabbit HQ following by DISCOVERY anti-HQ-HRP incubation. For double IHC stain, two antigens were sequentially detected and heat inactivation was used to prevent antibody cross-reactivity between the same species. Following each primary antibody incubation, DISCOVERY anti-Rabbit HQ or NP or DISCOVERY anti-Mouse HQ or NP and DISCOVERY anti-HQ-HRP or anti-NP-AP were incubated. The stains were visualized with DISCOVERY ChromoMap DAB Kit, DISCOVERY Yellow Kit, DISCOVERY Teal Kit or DISCOVERY Purple Kit; accordingly, counterstained with hematoxylin (Ventana) and coverslipped. The following primary Antibody information were listed: SPIKE (40150-T62, Sino Biological, at 1:2000), NP (NB100-56576, NOVUS, 1/100), hACE2 (AMAB91262, SIGMA, at 1:1000), CC10 (SC-365992#, Santa Cruz at 1/5000), Pro-SPC (AB37386, Millipore at 1/500), CD68 (ab125212, abcam, at 1/100) and NeuN (24307, cell signaling, at 1/100).

### Statistical analysis

Comparison of 2 groups was done using Student’s two-tailed t-test (for unpaired samples) or a paired t-test (for paired samples). Multiple groups were compared using one-way or two-way ANOVA and P values were adjusted for multiple comparisons by Holm’s procedure. A P value of 0.05 or less was considered statistically significant. Kaplan-Meier analysis was performed on the survival curves.

## Acknowledgement

Research reported in this publication included work performed in the Pathology Core supported by the National Cancer Institute of the National Institutes of Health under grant number P30CA033572. The content is solely the responsibility of the authors and does not necessarily represent the official views of the National Institutes.

## Author contributions

J. Yu, M.A. Caligiuri, P. Keim, B. Barker, H. Mead and W. Dong conceived and designed the project. P. Keim and B. Barker supervised experiments conducted in the laboratories. W. Dong, H. Mead, S. Jaramillo, D. Kollath, V. Coyne, N. Stone, A. Jones, J. Zhang and A. Li performed experiments and/or data analyses. M. Milanes-Yearsley reviewed the pathology changes. W. Dong, J. Yu, H. Mead, L-S Wang, and M.A. Caligiuri wrote, reviewed and/or revised the paper. All authors discussed the results and commented on the manuscript.

## Competing interest

The authors declare that they have no competing interests.

## Notes

### Competing Interest Statement

The authors have declared no competing interest.

## References

1. Davison AC, Hinkley DV. 1997. Bootstrap methods and their application. Cambridge University Press, Cambridge; New York, NY, USA.

2. Lauer SA, Grantz KH, Bi Q, Jones FK, Zheng Q, Meredith HR, Azman AS, Reich NG, Lessler J. 2020. The Incubation Period of Coronavirus Disease 2019 (COVID-19) From Publicly Reported Confirmed Cases: Estimation and Application. Ann Intern Med 172:577–582.

3. WHO. 2021. Report of the WHO-China Joint Mission on Coronavirus Disease 2019 (COVID-19)

4. Gibson PG, Qin L, Puah SH. 2020. COVID-19 acute respiratory distress syndrome (ARDS): clinical features and differences from typical pre-COVID-19 ARDS. Med J Aust 213:54–56 e1.

5. Brodin P. 2021. Immune determinants of COVID-19 disease presentation and severity. Nat Med 27:28–33.

6. Blot M, Bour JB, Quenot JP, Bourredjem A, Nguyen M, Guy J, Monier S, Georges M, Large A, Dargent A, Guilhem A, Mouries-Martin S, Barben J, Bouhemad B, Charles PE, Chavanet P, Binquet C, Piroth L, group Ls. 2020. The dysregulated innate immune response in severe COVID-19 pneumonia that could drive poorer outcome. J Transl Med 18:457.

7. Mazzoni A, Salvati L, Maggi L, Capone M, Vanni A, Spinicci M, Mencarini J, Caporale R, Peruzzi B, Antonelli A, Trotta M, Zammarchi L, Ciani L, Gori L, Lazzeri C, Matucci A, Vultaggio A, Rossi O, Almerigogna F, Parronchi P, Fontanari P, Lavorini F, Peris A, Rossolini GM, Bartoloni A, Romagnani S, Liotta F, Annunziato F, Cosmi L. 2020. Impaired immune cell cytotoxicity in severe COVID-19 is IL-6 dependent. J Clin Invest 130:4694–4703.

8. Walls AC, Park YJ, Tortorici MA, Wall A, McGuire AT, Veesler D. 2020. Structure, Function, and Antigenicity of the SARS-CoV-2 Spike Glycoprotein. Cell 181:281–292 e6.

9. McCray PB, Jr., Pewe L, Wohlford-Lenane C, Hickey M, Manzel L, Shi L, Netland J, Jia HP, Halabi C, Sigmund CD, Meyerholz DK, Kirby P, Look DC, Perlman S. 2007. Lethal infection of K18-hACE2 mice infected with severe acute respiratory syndrome coronavirus. J Virol 81:813–21.

10. Jiang RD, Liu MQ, Chen Y, Shan C, Zhou YW, Shen XR, Li Q, Zhang L, Zhu Y, Si HR, Wang Q, Min J, Wang X, Zhang W, Li B, Zhang HJ, Baric RS, Zhou P, Yang XL, Shi ZL. 2020. Pathogenesis of SARS-CoV-2 in Transgenic Mice Expressing Human Angiotensin-Converting Enzyme 2. Cell 182:50–58 e8.

11. Bao L, Deng W, Huang B, Gao H, Liu J, Ren L, Wei Q, Yu P, Xu Y, Qi F, Qu Y, Li F, Lv Q, Wang W, Xue J, Gong S, Liu M, Wang G, Wang S, Song Z, Zhao L, Liu P, Zhao L, Ye F, Wang H, Zhou W, Zhu N, Zhen W, Yu H, Zhang X, Guo L, Chen L, Wang C, Wang Y, Wang X, Xiao Y, Sun Q, Liu H, Zhu F, Ma C, Yan L, Yang M, Han J, Xu W, Tan W, Peng X, Jin Q, Wu G, Qin C. 2020. The pathogenicity of SARS-CoV-2 in hACE2 transgenic mice. Nature 583:830–833.

12. Sun SH, Chen Q, Gu HJ, Yang G, Wang YX, Huang XY, Liu SS, Zhang NN, Li XF, Xiong R, Guo Y, Deng YQ, Huang WJ, Liu Q, Liu QM, Shen YL, Zhou Y, Yang X, Zhao TY, Fan CF, Zhou YS, Qin CF, Wang YC. 2020. A Mouse Model of SARS-CoV-2 Infection and Pathogenesis. Cell Host Microbe 28:124–133 e4.

13. Hassan AO, Case JB, Winkler ES, Thackray LB, Kafai NM, Bailey AL, McCune BT, Fox JM, Chen RE, Alsoussi WB, Turner JS, Schmitz AJ, Lei T, Shrihari S, Keeler SP, Fremont DH, Greco S, McCray PB, Jr., Perlman S, Holtzman MJ, Ellebedy AH, Diamond MS. 2020. A SARS-CoV-2 Infection Model in Mice Demonstrates Protection by Neutralizing Antibodies. Cell doi:10.1016/j.cell.2020.06.011.

14. Grant RA, Morales-Nebreda L, Markov NS, Swaminathan S, Querrey M, Guzman ER, Abbott DA, Donnelly HK, Donayre A, Goldberg IA, Klug ZM, Borkowski N, Lu Z, Kihshen H, Politanska Y, Sichizya L, Kang M, Shilatifard A, Qi C, Lomasney JW, Argento AC, Kruser JM, Malsin ES, Pickens CO, Smith SB, Walter JM, Pawlowski AE, Schneider D, Nannapaneni P, Abdala-Valencia H, Bharat A, Gottardi CJ, Budinger GRS, Misharin AV, Singer BD, Wunderink RG, Investigators NSS. 2021. Circuits between infected macrophages and T cells in SARS-CoV-2 pneumonia. Nature 590:635–641.

15. Chow YH, O’Brodovich H, Plumb J, Wen Y, Sohn KJ, Lu Z, Zhang F, Lukacs GL, Tanswell AK, Hui CC, Buchwald M, Hu J. 1997. Development of an epithelium-specific expression cassette with human DNA regulatory elements for transgene expression in lung airways. Proc Natl Acad Sci U S A 94:14695–700.

16. Yang XH, Deng W, Tong Z, Liu YX, Zhang LF, Zhu H, Gao H, Huang L, Liu YL, Ma CM, Xu YF, Ding MX, Deng HK, Qin C. 2007. Mice transgenic for human angiotensin-converting enzyme 2 provide a model for SARS coronavirus infection. Comp Med 57:450–9.

17. Ostrowski LE, Hutchins JR, Zakel K, O’Neal WK. 2003. Targeting expression of a transgene to the airway surface epithelium using a ciliated cell-specific promoter. Mol Ther 8:637–45.

18. Arce VM, Costoya JA. 2021. SARS-CoV-2 infection in K18-ACE2 transgenic mice replicates human pulmonary disease in COVID-19. Cell Mol Immunol 18:513–514.

19. Moreau GB, Burgess SL, Sturek JM, Donlan AN, Petri WA, Mann BJ. 2020. Evaluation of K18-hACE2 Mice as a Model of SARS-CoV-2 Infection. Am J Trop Med Hyg 103:1215–1219.

20. Oladunni FS, Park JG, Pino PA, Gonzalez O, Akhter A, Allue-Guardia A, Olmo-Fontanez A, Gautam S, Garcia-Vilanova A, Ye C, Chiem K, Headley C, Dwivedi V, Parodi LM, Alfson KJ, Staples HM, Schami A, Garcia JI, Whigham A, Platt RN, 2nd, Gazi M, Martinez J, Chuba C, Earley S, Rodriguez OH, Mdaki SD, Kavelish KN, Escalona R, Hallam CRA, Christie C, Patterson JL, Anderson TJC, Carrion R, Jr., Dick EJ, Jr., Hall-Ursone S, Schlesinger LS, Alvarez X, Kaushal D, Giavedoni LD, Turner J, Martinez-Sobrido L, Torrelles JB. 2020. Lethality of SARS-CoV-2 infection in K18 human angiotensin-converting enzyme 2 transgenic mice. Nat Commun 11:6122.

21. Winkler ES, Bailey AL, Kafai NM, Nair S, McCune BT, Yu J, Fox JM, Chen RE, Earnest JT, Keeler SP, Ritter JH, Kang LI, Dort S, Robichaud A, Head R, Holtzman MJ, Diamond MS. 2020. SARS-CoV-2 infection of human ACE2-transgenic mice causes severe lung inflammation and impaired function. Nat Immunol 21:1327–1335.

22. Yinda CK, Port JR, Bushmaker T, Owusu IO, Avanzato VA, Fischer RJ, Schulz JE, Holbrook MG, Hebner MJ, Rosenke R, Thomas T, Marzi A, Best SM, de Wit E, Shaia C, van Doremalen N, Munster VJ. 2020. K18-hACE2 mice develop respiratory disease resembling severe COVID-19. bioRxiv doi:10.1101/2020.08.11.246314.

23. Sia SF, Yan LM, Chin AWH, Fung K, Choy KT, Wong AYL, Kaewpreedee P, Perera R, Poon LLM, Nicholls JM, Peiris M, Yen HL. 2020. Pathogenesis and transmission of SARS-CoV-2 in golden hamsters. Nature 583:834–838.

24. Sepehrinezhad A, Shahbazi A, Negah SS. 2020. COVID-19 virus may have neuroinvasive potential and cause neurological complications: a perspective review. J Neurovirol 26:324–329.

25. Muhammad S, Petridis A, Cornelius JF, Hanggi D. 2020. Letter to editor: Severe brain haemorrhage and concomitant COVID-19 Infection: A neurovascular complication of COVID-19. Brain Behav Immun 87:150–151.

26. Mao XY, Jin WL. 2020. The COVID-19 Pandemic: Consideration for Brain Infection. Neuroscience 437:130–131.

27. Ramani A, Pranty AI, Gopalakrishnan J. 2021. Neurotropic effects of SARS-CoV-2 modeled by the human brain organoids. Stem Cell Reports doi:10.1016/j.stemcr.2021.02.007.

28. Pellegrini L, Albecka A, Mallery DL, Kellner MJ, Paul D, Carter AP, James LC, Lancaster MA. 2020. SARS-CoV-2 Infects the Brain Choroid Plexus and Disrupts the Blood-CSF Barrier in Human Brain Organoids. Cell Stem Cell 27:951–961 e5.

29. Zhang BZ, Chu H, Han S, Shuai H, Deng J, Hu YF, Gong HR, Lee AC, Zou Z, Yau T, Wu W, Hung IF, Chan JF, Yuen KY, Huang JD. 2020. SARS-CoV-2 infects human neural progenitor cells and brain organoids. Cell Res 30:928–931.

30. Cooper KW, Brann DH, Farruggia MC, Bhutani S, Pellegrino R, Tsukahara T, Weinreb C, Joseph PV, Larson ED, Parma V, Albers MW, Barlow LA, Datta SR, Di Pizio A. 2020. COVID-19 and the Chemical Senses: Supporting Players Take Center Stage. Neuron 107:219–233.

31. Stone NE, Jaramillo SA, Jones AN, Vazquez AJ, Martz M, Versluis LM, Raniere MO, Nunnally HE, Zarn KE, Nottingham R, Ng KR, Sahl JW, Wagner DM, Knudsen S, Settles EW, Keim P, French CT. 2021. Stenoparib, an Inhibitor of Cellular Poly(ADP-Ribose) Polymerase, Blocks Replication of the SARS-CoV-2 and HCoV-NL63 Human Coronaviruses In Vitro. mBio 12.

